# Semi-automated navigation for efficient targeting of electron tomography to regions of interest in volume correlative light and electron microscopy

**DOI:** 10.1101/2024.11.29.626074

**Authors:** Kohki Konishi, Guilherme Neves, Matthew Russell, Masafumi Mimura, Juan Burrone, Roland Fleck

## Abstract

Electron microscopy is essential for the quantitative study of synaptic ultrastructure. At present, the correlation of functional and structural properties of the same synapse is extremely challenging. We introduce a novel integrated workflow designed to simplify sample navigation across spatial scales, allowing the identification of individual synapses from optical microscopy mouse brain image stacks that can be targeted for analysis using electron tomography imaging. We developed a software which has a function to register multimodal images using a novel segmentation-based image registration algorithm as well as a function to visualize all the registration results. Using our newly designed software we streamline mapping of high-resolution optical imaging onto reference maps using blood vessels as endogenous fiducial marks. Further we demonstrate significant improvements on the ultramicrotomy stage of volume Correlative Light and Electron Microscopy (vCLEM) workflows, providing real time guidance to targeted trimming to match previously acquired Regions Of Interest (ROIs), and reliable estimates of cutting depth relative to ROI, based on fluorescence imaging of TEM ready ultrathin sections. Using this workflow, we successfully targeted TEM tomography to the proximal axonal region containing the Axon Initial Segment identified using fluorescent light microscopy.

## Introduction

Electron microscopy (EM) has been a powerful tool for investigating the ultrastructure of cells and tissues. Volume EM (vEM), which enables three-dimensional (3D) ultrastructural analysis, has gained significant attention [1,2]. Block-face methods, such as serial block-face scanning electron microscopy (SBF-SEM; [3]) and focused ion beam scanning electron microscopy (FIB-SEM; [4]), are widely used for acquiring vEM data by sequentially imaging the surface of a sample block. SBF-SEM has yielded novel insights into development of synaptic ultrastructure [5,6]. Other volume methods include serial section electron microscopy (ssTEM), where serial sections are collected on grids and imaged using transmission electron microscopy (TEM; [7]). ssTEM and SBF-SEM have been extensively used in connectomics research to analyze neural circuit connectivity (e.g. [8,9]).

Volume correlative light and electron microscopy (vCLEM) enables the visualization of the ultrastructure of specific targets with known functional properties by volume imaging of fluorescently labeled cells or tissues (e.g. [10] reviewed by [11–13]). Recently, A workflow that performs multiplexed detergent-free immunolabeling and vCLEM on the same sample was developed by using mouse brain tissue [14]. However, navigation to the target of interest across different microscopy modalities and different scales remains labor-intensive.

Image registration is a crucial technique for navigation between microscopy images of the same structures across scales and imaging modalities. At each stage, alignment of an image from one modality and/or scale with another is needed so that one can be used as a reference to navigate to the same region of interest within the other. A range of registration approaches have been developed. One conventional method is for a user to manually identify landmark features that are visible in both images (eg cell nuclei). A transform of one image – rotation, translation, non-linear warping, etc. – can then be calculated so that corresponding landmark features are aligned when the image is overlaid onto the other one. This approach is enabled by tools such as Bigwarp [15] and ec-CLEM [16]. While these methods can achieve high accuracy, manual landmark selection process is labor-intensive. Consequently, automating multimodal image registration has been investigated. AutoCLEM was developed to automatically detect embedded beads and use them as landmarks for image warping [17]. Intensity-based image registration tools are freely provided as the Elastix toolbox [18,19] for medical image registration, implemented in a multi-modal big image data sharing and exploration viewer (MoBIE; [20]), and applied to align DAPI signals with segmented nuclear masks from EM images, enabling the identification of cells corresponding to gene expression [21]. Moreover, combination of segmentation and point cloud matching was developed to automatically establish correspondences between two images [22].

Recent years have seen a surge in research on efficient navigation workflows to fluorescently labeled targets for vEM. One approach involves acquiring light microscopy images of fluorescently labeled cells, followed by resin embedding and vEM imaging using the ‘approach and correlation’ method [23]. Another workflow embeds the sample in resin, then iteratively performs light microscopy imaging and stepwise trimming to navigate to fluorescently labeled cells for FIB-SEM imaging [24,25]. A third approach incorporates x-ray micro-CT imaging between light microscopy and vEM imaging [26–29]. Additionally, a workflow integrating a motorized ultramicrotome with x-ray micro-CT imaging has been proposed [30]. This approach also requires access to x-ray micro-CT.

In this paper we present a novel integrated workflow for vCLEM that uses in resin fluorescence signals to allow navigation of millimetre sized resin embedded mouse brain tissue blocks and specific targeting of sub-micrometre sized regions containing synaptic structures for TEM tomography. Our strategy starts with the creation of a reference map using fluorescence microscopy confocal stacks encompassing the entire tissue block. Regions Of Interest (ROIs) are then imaged at higher resolution and mapped onto the reference map using a newly created user-friendly software based on napari (NavROI) that simplifies registering and display of imaging datasets acquired with different imaging modalities. Previously acquired confocal images can be registered in real time with the video signal of the ultramicrotome camera to simplify the process of sample trimming in the X-Y dimension. Further, we show that the fluorescence of ultrathin (200 nm) sections can be mapped with the reference to produce reliable estimates of section Z depth. We used this approach to target TEM tomography to the proximal axonal region containing the Axon Initial Segment fluorescently labelled with a viral approach.

## Results

### Integrated workflow

We developed an integrated workflow of navigating ROIs for volume CLEM. Figure 1(A) shows its overview. The workflow integrates imaging and computation across spatial scales, from light microscopy (LM) imaging of sub-millimeter scale to highlighting individual subcellular structure, to transmission electron microscopy (TEM) tomogram imaging of nanometer scale structures. The computation consists of two algorithms to register pairs of images in different modalities and/or scales, to facilitate navigation through them. The two algorithms are, 1) a novel segmentation-based image registration algorithm shown in Figure 1(B), which is described in the next section, and 2) a conventional landmark-based image registration algorithm.

**Figure 1:**
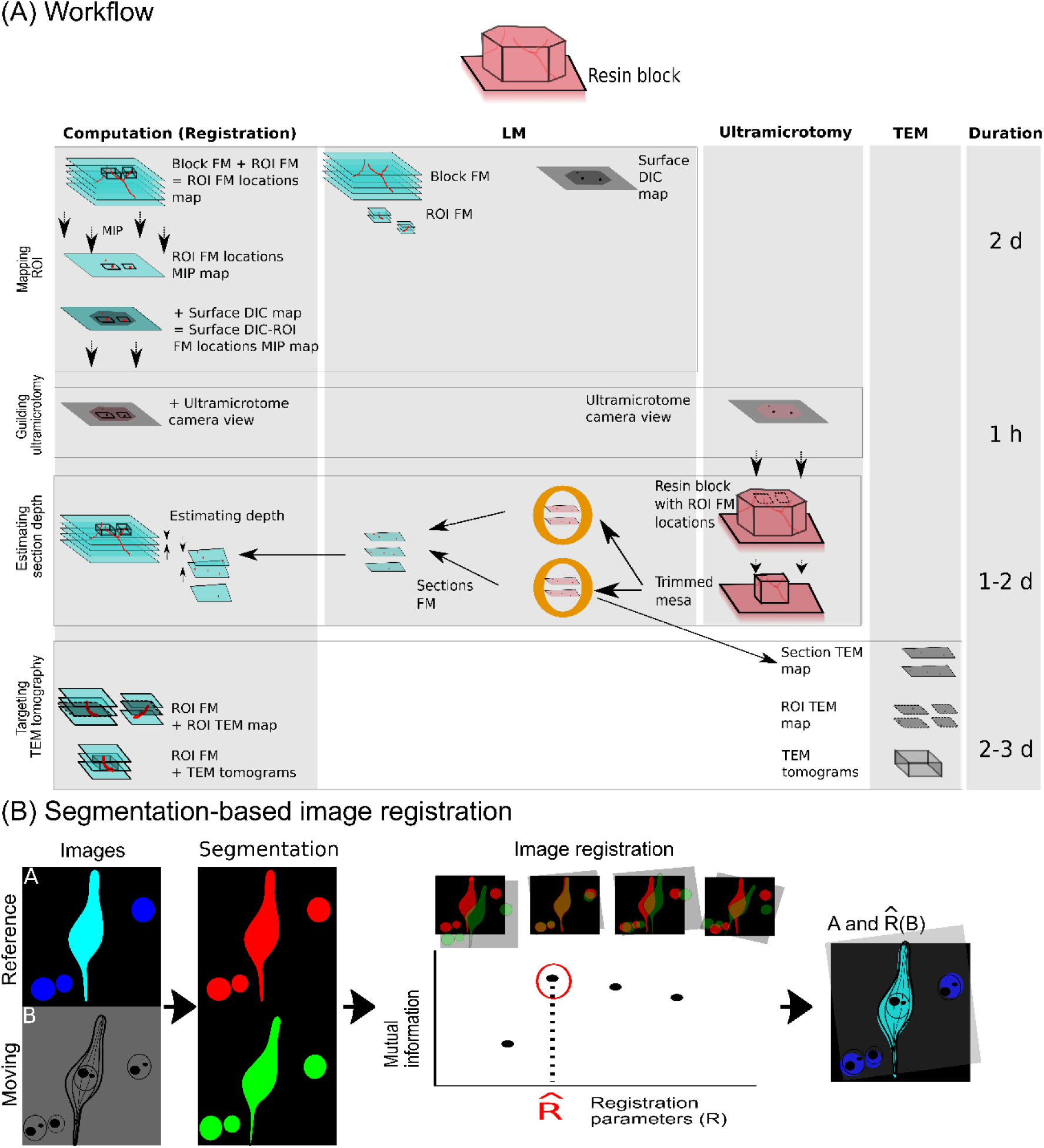
Integrated workflow. (A) Integrated workflow which integrates imaging and computation. Imaging consists of light microscopy (LM) imaging, ultramicrotomy, transmission electron microscopy (TEM). The computation consists of two image registration algorithms: segmentation-based image registration, shown in panel (B), and conventional landmark-based image registration. The workflow comprises four steps: mapping ROI (duration, 2 days), guiding ultramicrotomy (1 hour), estimating section depth (1-2 days) and targeting TEM tomography (2-3 days). If all the steps are performed without interruption, the average duration of this workflow is thus roughly 1 week. (B) Segmentation-based image registration algorithm. The algorithm consists of two primary components: segmentation and registration. The segmentation step segments common objects in the images from both modalities, either automatically or with user assistance. The registration step maximizes a metric called mutual information (MI) while warping the moving image. See the text for details.

The workflow starts with a resin block containing brain tissue with fluorescently tagged markers (Figure 1(A)). We initially identified regions of interest (ROIs) within the block by microscopy of the fluorescent markers, and trimmed the sides of the block to those ROIs. To direct this trimming process we needed to be able to identify the lateral position of those regions when viewing the block surface in the ultramicrotome. We therefore registered pairs of images that could be used to navigate between the ROIs within the block and the view of the full surface of the block.

We first acquired a full-width fluorescence microscopy (FM) image stack of the resin block (Block FM, Figure 1). The regions of interest (ROIs) were visually identified and subsequently imaged at higher resolution (ROI FM, Figure 1). To spatially register these image datasets, we generated a three-dimensional map of the ROI locations within the entire block through segmentation-based image registration (ROI FM locations map, Figure 1). Since FM features within the block are not visible in the ultramicrotome stereomicroscope, a DIC image of the full surface of the resin block was also acquired at this point to relate the FM ROIs to the surface view in the ultramicrotome (Surface DIC map, Figure 1). A maximum intensity projection (MIP) of the ROI FM locations map (ROI FM locations MIP map, Figure 1) was then generated. Landmark-based image registration was then employed to register the ROI FM locations MIP map with the Surface DIC map, essentially resulting in a projection of the lateral positions of the ROIs onto the surface view of the block (Surface DIC-ROI FM locations MIP map, Figure 1).

The subsequent step involved guiding the lateral trimming of the block during ultramicrotomy. The ultramicrotome camera view of the block was acquired and registered with the Surface DIC-ROI FM locations MIP map through landmark-based image registration. This process enabled the overlay of the projected lateral ROI locations onto the ultramicrotome camera view. By referencing this overlay during the trimming process, we optimized mesa trimming by removing material outside the ROI FM.

We then proceeded with sectioning into the trimmed mesa to find sections at the depth of the ROI FMs. To find the sections at the correct depth, we collected sections onto cover slips and imaged using FM (Sections FM, in Figure 1). We defined the depth of each section as the depth corresponding to the optimal segmentation-based image registration between Section FM and Block FM. By this method we could estimate the depth of each section within the Block FM, and then relate this to the ROI FM locations. Thus, the sections whose estimated cutting depth matches the depth of the ROI FMs in the ROI FM locations map could be selected for subsequent correlative TEM imaging of those ROIs.

The final step involved targeting TEM tomography to specific ROIs identified in the high resolution ROI FM image stacks. TEM images of the whole of selected sections (Section TEM map, Figure 1) were acquired. Within these images, specific ROIs were located and targeted for higher magnification TEM imaging (ROI TEM map, Figure 1). Segmentation-based image registration was employed to register the ROI TEM map with the ROI FM map, enabling the overlay of target positions onto the ROI TEM map. TEM tomography was then performed at the target positions. Through the same image registration method, the ROI FM was successfully registered with the TEM tomogram. If all the steps are performed without interruption, the average duration of this workflow is roughly 1 week.

### Segmentation-based image registration for multimodal images

We developed a segmentation-based image registration algorithm for registering pairs of multimodal images such as images acquired from different microscopy modalities and/or scales. Figure 1(B) presents a flowchart illustrating the algorithm. One image serves as the reference image (Image A in Figure 1B), while the other serves as the moving image (Image B in Figure 1B). The algorithm primarily consists of two primary components: object segmentation and image registration.

For object segmentation, we employed automated or user-assisted techniques to segment objects that are frequently present in both images. Segmentation masks were subsequently converted into labels via connected component labeling. Examples of such common objects include blood vessels, cell nuclei, and neurons.

During image registration, we selected the registration that yielded the highest mutual information (MI) metric. To achieve this, we randomly generated affine transformation parameters, warped the moving image, and computed the MI values between the labels from images A and B.

### ROI navigation tool

To streamline the integrated workflow, we developed a ROI navigation tool, navROI, using napari that displays the spatial relationships between multiple images. The images can be multiple image stacks, multiple section images, or their mixture. The tool accepts the paths of the images and their registration parameters and displays their locations relative to the largest field-of-view image.

### Implementation of the integrated workflow to mouse brain tissue

We applied the integrated workflow to a resin block in which mouse brain tissue was embedded.

#### Mapping the ROI FM to the Block FM

Figure 2(A) shows an example of ROI FMs of mouse brain tissue. Neurons are shown in cyan, synapses in green, blood vessels in red, and nuclei in blue. The neuron centered in the ROI FM is surrounded by distinctive blood vessels. Three points of the blood vessel structures are indicated by five arrowheads. Since the blood vessel structure is diverse, we used the blood vessel channel to map the ROI FM locations to the Block FM. The region containing the ROI FM is highlighted in the white box and its volume rendering is shown in the inset. The five green arrowheads in panel (B) are the corresponding points in panel (A).

**Figure 2:**
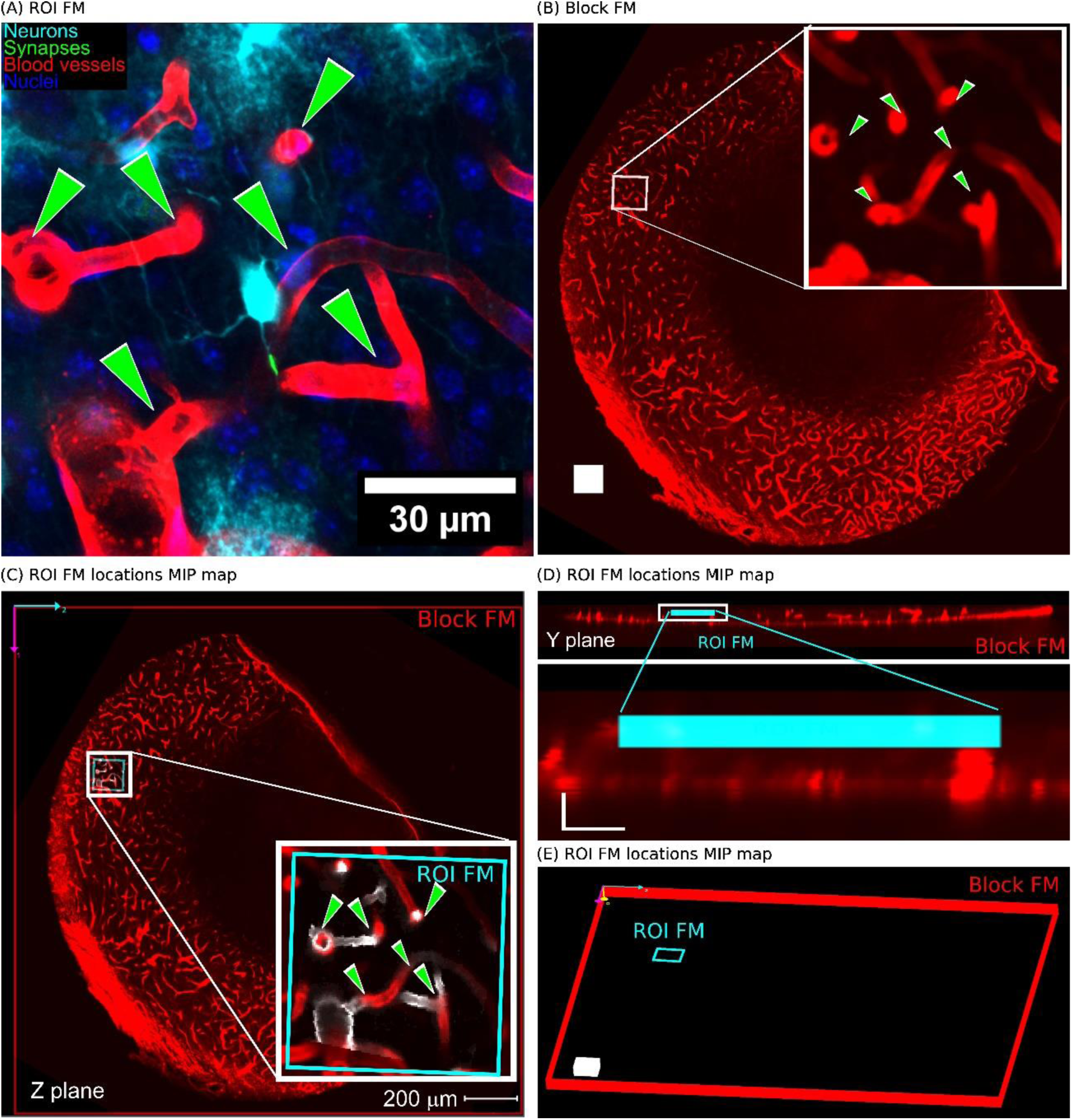
Mapping the ROI FM within the Block FM. (A) ROI FM of mouse brain tissue: Neurons in cyan, synapses in green, blood vessels in red, and nuclei in blue are visualized. Three structures in the blood vessel channel are indicated by green arrowheads. (B) Block FM blood vessel channel: A volume rendering highlights the region containing the ROI FM, white box. A zoomed-in view is shown in the inset, with five structures corresponding to those in panel a marked by green arrowheads. The size of white cube is 100 microns. (C)-(E) ROI FM Location Maps: (C) navROI Z-plane view: A screenshot of the navROI tool in Z-plane view. Blood vessels from ROI FM are shown in white. (D) navROI Y-plane view: Screenshots of navROI in Y-plane view, showing both the entire view, top panel, and a zoomed-in region around the ROI FM, bottom panel. The size of the L is 20 microns. (E) Spatial relationship: A navROI screenshot illustrating the spatial relationship between two regions of the ROI FM and the Block FM.

To map the ROI FM to the Block FM, we utilized image registration and visualization techniques. Segmentation-based image registration was employed to determine the optimal registration parameters between these two image stacks. Subsequently, we utilized navROI for visualization by inputting the file paths of the ROI FM, Block FM, and the optimal registration parameters.

Figures 2(C) and 2(D) present screenshots of the navROI interface. Figure 2(C) displays its Z-plane view, where the ROI FM region is highlighted with a cyan-colored box, the neuron channel of the ROI FM is visualized, and the blood vessel channel of the Block FM is also visualized in red. A zoomed-in view of the white box surrounding the ROI FM is shown in the lower right corner. Figure 2(D) illustrates the Y-plane view of the navROI, with the upper panel showing an overall view and the lower panel showing a zoomed-in view of the white box around the ROI FM.

Figure 2(E) depicts a screenshot of a bird’s-eye view generated by navROI, enabling visualization of the mapping the ROI FM region in cyan to the Block FM region in red. This visualization is utilized for lateral trimming of blocks and section selection, as detailed below.

#### Guiding the lateral trimming of the block during ultramicrotomy

We optimized trimming of the block mesa by overlaying ROI FM locations onto the ultramicrotome (UC) view of the block. Figure 3(A) illustrates a schematic of the UC system, with the camera view displayed on the top screen. Figure 3(B) presents a UC camera view of the block before trimming. Three landmarks of the block corners are marked with blue arrowheads. Figure 3(C) shows a Surface DIC map of the block. The landmarks corresponding to those in panel (B) are indicated by blue arrowheads, while three blood vessel landmarks are marked with red arrowheads. Figure 3(D) displays a ROI locations MIP map. The five ROI locations are highlighted as green rectangles, and the three landmarks of the blood vessel corner locations corresponding to those in panel (C) are marked with red arrowheads. Since the surface DIC map is registered with the ROI FM locations MIP map, we obtained an overlay of ROI locations on the UC camera view. Figure 3(E) shows the overlay where the trimming region enclosing all the ROIs is highlighted in pink. We trimmed away materials outside of the pink polygonal region using this overlay as a guide. Figure 3(F) shows an UC camera view of the trimmed block.

**Figure 3:**
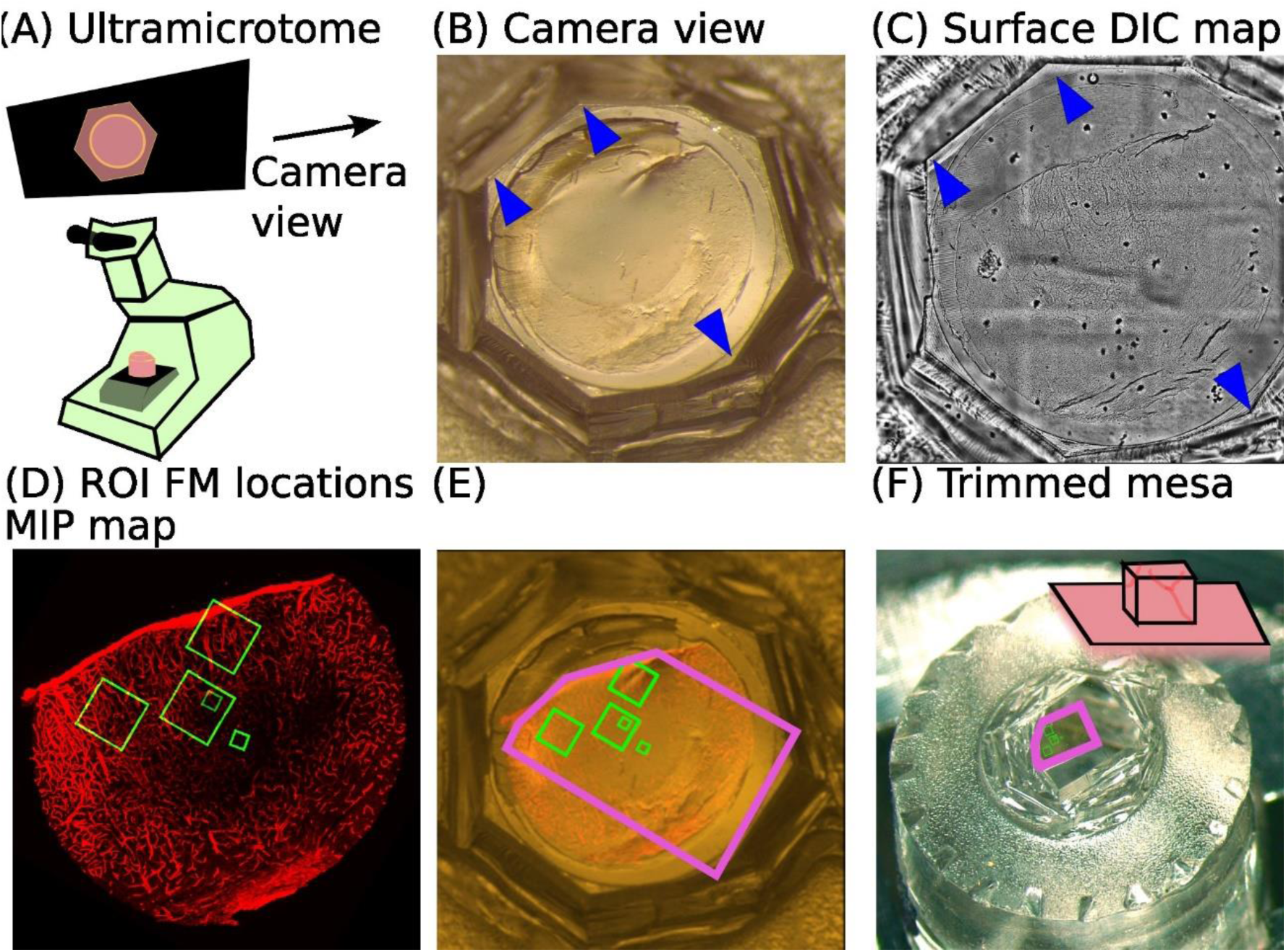
Using a Surface Differential Interference Contrast (DIC) map of the block to guide ultramicrotomy. (A) A schematic of the UC and its camera view on top. (B) An UC camera view of the block. Three landmarks of the block corners are indicated by blue arrowheads. (C) A Surface DIC map of the block. The landmarks corresponding to those in panel b are indicated by blue arrowheads, and three landmarks of the blood vessels in red. (D) A ROI FM locations MIP map. Five regions of ROI FMs are shown in green. (E) A screen shot of navROI illustrating an UC camera view overlaid with five ROI FMs. The trimming region is shown in pink. The Surface DIC map and the ROI FM locations MIP map are also shown. (F) An UC camera view of the trimmed block. The trimmed region is shown in pink.

#### Estimating cutting depth of sections

We estimated cutting depth of sections by reading the depth from the Block FM surface at the point of optimal image registration between Section FM and Block FM through segmentation-based image registration. Figure 4(A) shows a local region of blood vessel channel image of Section FM, with the full section shown in the inset. We focused image registration analysis to the local region of a few hundred-micron size to mitigate non-linear distortions in the Section FM. Subsequently, we utilized navROI for the mapping process by inputting the file paths of the entire Section FM, its local region, Block FM, and the optimal registration parameters.

**Figure 4:**
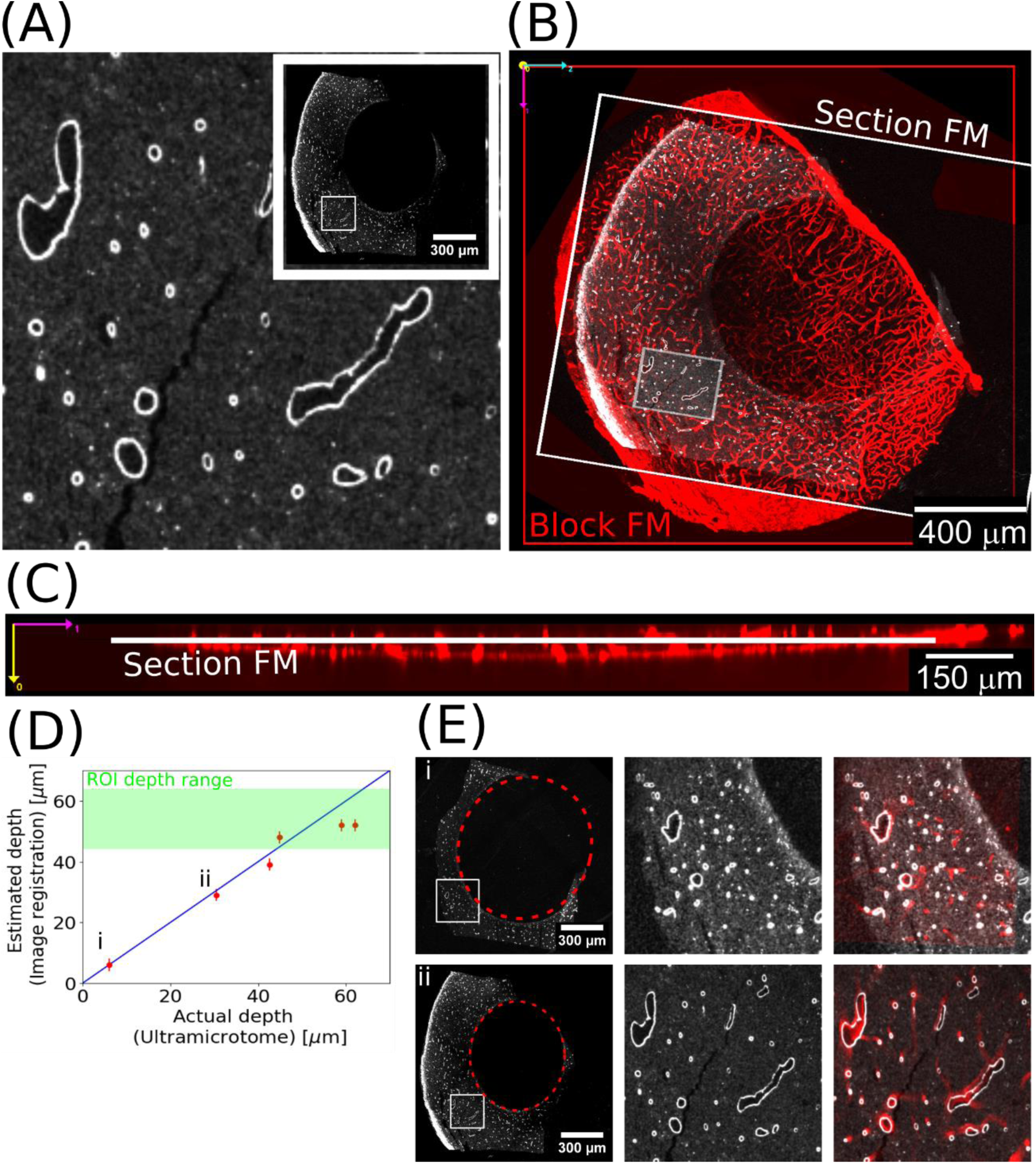
Estimating cutting depth of sections. (A) A local region of the blood vessel channel image of a Section FM. Its entire image is shown in the inset. (B)-(C) A Section FM is displayed relative to the Block FM using navROI: (B) A screen shot of navROI in Z plane view. (C) A screen shot of navROI in Y plane view. (D) Comparison of the feeding values from the UC and the depth estimation using image registration. The details of depth estimation for the leftmost point *i* and point *ii* to its right in the plot are shown in panel E. (E) The image registration for depth estimation. Left: An entire section image with a white rectangle. The white rectangle region is used for depth estimation. The inside the dotted red circle shows an empty resin region. Middle: The zoomed-in view of the white rectangle region. Right: The result of image registration for depth estimation. The top row: point *i*, the bottom row: point *ii*.

Figures 4(B) and 4(C) present screenshots of the navROI interface. Figure 4(B) displays its Z-plane view, where the Section FM and its local region are highlighted with white boxes, and the blood vessel channel of the Block FM is visualized in red. Figure 4(C) illustrates the Y-plane view of the navROI. This view visualizes the depth of the Section FM from the surface of the Block FM.

Figure 4(D) compares the cutting depths of the six sections relative to the block surface with the corresponding feeding values from the UC. Section FMs were used to generate first two points nearest to the block surface and Section TEM maps were used to generate the remaining points. The green region indicates the depth range estimated from registering the ROI FM in the Block FM. A diagonal line with a slope of 1 is also plotted for reference. Linear regression analysis of the plot yielded a slope of 0.85 and an intercept of 3.00 micron. The first two sections closest to the block surface are labeled i and ii in the plot.

Figure 4(E) provides detailed process of the image registration for the Section FM labeled i and ii. The left panel displays the entire Section FM, with the local region used for registration highlighted in white rectangles. Note that the center of the Section FM contains no resin, as indicated by the dotted red circles. The middle panel shows a zoomed-in view of the local region in the left panel. This local region was utilized for image registration. The right panel presents the optimal image registration result between the Section FM and the Block FM. The section image is depicted in gray, while the corresponding region of the Block FM is shown in red.

### Targeting TEM tomography to specific ROIs in FM images

In the previous sections, we utilized blood vessels in navigating ROIs. Here, we show navigation from the FM ROI to targeted TEM tomography of targets along an axon identified within the FM ROI using nuclei and neurons.

Figure 5(A) presents a Section TEM map with an overlaid ROI FM. The ROI FM is marked by a cyan box. A neuron is shown in cyan, and nuclei in blue. A white box emphasizes a region containing the axon, which is encircled by three nuclei indicated by yellow arrowheads.

**Figure 5:**
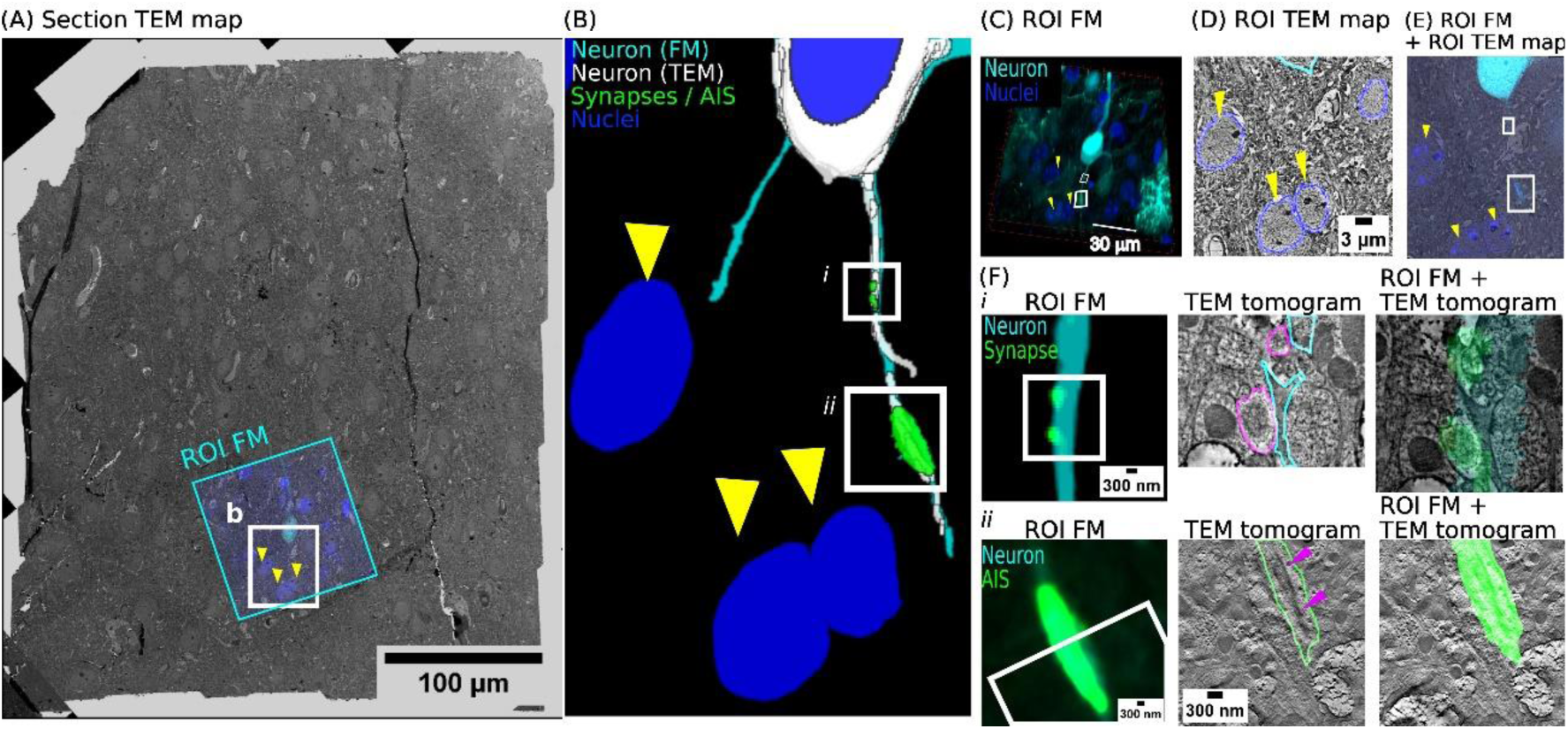
Targeting TEM tomography to synapses identified in confocal images. (A) An overlay of a ROI FM on a Section TEM map. The region surrounded by a white rectangle contains an axon of a neuron and is magnified on the right panel. Surrounding nuclei are indicated by yellow arrowheads. (B) A volume rendering of the segmentations of the neuron from the ROI FM and the ROI TEM map, displayed in cyan and white. The two target regions for TEM tomography are shown in green and highlighted in a white rectangle. Surrounding nuclei are shown in blue. (C) A three-dimensional view of the ROI FM. (D) An example of the ROI TEM map with segmentation. (E) The overlay of the ROI FM on the ROI TEM map. (F) Image registration at the target regions. Left: The target region in the ROI FM, the cytoplasmic marker channel in cyan and synapse channel in green. The corresponding region to the Middle panel is shown in white rectangle. Middle: The TEM tomogram around the region with segmentation. Right: Their overlay. The top row: *i*. Synapses in the axon. Presynaptic boutons, outlined in purple, correspond to postsynaptic densities identified by fluorescent labeling. The bottom row: *ii*. A blob in the axon initial segment (AIS). Cisternal organelle, indicated by a purple arrowhead, is a distinctive feature expected to be found in the AIS.

Figure 5(B) presents a volume rendering of the axon segmented from the ROI FM and the ROI TEM map, shown in cyan and white, respectively. Segmentation of synapses along the axon of the neuron and a blob in the axon initial segment (AIS) from the ROI FM are shown in green. These are the TEM tomography targets and are further emphasized by white rectangles labeled i or ii. Additionally, nuclei segmentation from the ROI FM are rendered in blue with yellow arrowheads.

To reconstruct the axon from serial ROI TEM maps, these maps should be registered. We registered each ROI TEM map to the FM Block using segmentation-based image registration algorithm. Figure 5(C) presents a volume rendering of the FM Block, highlighting neurons and nuclei in cyan and blue. The two target regions are outlined by white rectangles. Figure 5(D) shows an example ROI TEM map, overlaying neuron and nuclei segmentations in cyan and blue. Figure 5(E) demonstrates optimal image registration between panels (C) and (D). By examining Figure 5(E), we segmented the axon from the ROI TEM map. Repeating this registration process for all ROI TEM maps achieved axon segmentation and subsequent reconstruction.

To identify targets in ROI TEM maps, we performed non-linear image registration between adjacent ROI TEM maps by focusing on the local region of a few ten-micron size containing targets. This additional step was necessary because the initial linear image registration was not sufficient to trace the axon across adjacent ROI TEM maps due to residual non-linear registration errors. By employing non-linear image registration, we overlaid the fluorescent signal of targets onto the ROI TEM maps. After non-linear image registration, we proceeded to acquire TEM tomograms.

We registered the FM Block to TEM tomograms through segmentation-based image registration as illustrated in Figure 5(F). Zoomed-in regions of ROI FM are shown in the left panel, alongside corresponding TEM tomograms with segmentation overlays, the middle panel, and optimal image registrations, the right panel. Upper panels depict the synapses in the axon, while lower panels show the AIS blob.

## Discussion

### Efficient navigation of ROIs using integrated vCLEM workflow

vCLEM offers unparalleled potential for exploring biological structures. Image registration techniques between light and electron microscopy images have been developed by several authors [15,16] and a central challenge in vCLEM has been the efficient navigation of ROIs across multimodal and multiscale images. Here we report an integrated vCLEM workflow with a novel image registration algorithm as well as a napari-based user-friendly visualization tool that simplifies the navigation from confocal microscopy images of the tissue block to high-resolution TEM tomography of the tissue sections. Our integrated workflow significantly improves both of robustness and efficiency of ROI navigation for vCLEM analysis.

### Robust, real-time ROI navigation with segmentation-based image registration

Robust and real-time image registration algorithms are crucial for ROI navigation. Conventional methods demand manual landmark correspondence between reference and moving images, a time-consuming task. To streamline this process, automated registration methods have been developed [17,22,31]. However, these methods often assume that both images are acquired from the same region, which may not be feasible in scenarios like serial section high-resolution TEM imaging.

Our novel image registration algorithm addresses this limitation by enabling immediate registration after acquiring a single image of a section, eliminating the need to wait for the entire series. By leveraging GPU acceleration for efficient mutual information (MI) computation, we achieve real-time performance. This allows for daytime imaging and same-night registration, providing timely feedback for informed decision-making. The next morning, users can decide whether to proceed with the planned imaging depth or adjust the number of sections to reach the desired ROI. This real-time capability enhances the robustness and efficiency of ROI selection. While we employed segmentation for image registration, generative adversarial network-based approaches [32] might also be applied to multi-modal image stack registration.

Multi-modal image registration algorithms have been extensively studied in medical imaging. For images where intensity values are positively correlated, such as MRI and CT images, MI has proven effective [33,34]. Segmentation-based approaches have been proposed for registering T1 and T2 MRI images [35]. Algorithms for image stack registration have also been actively explored [36].

Inspired by advancements in medical imaging, we propose a segmentation-based image registration method for vCLEM. Given the extensive research on segmenting light microscopy and TEM images [37], our method leverages these developments. We focused on segmenting key organelles: blood vessels, cell nuclei, and neurons. Blood vessel segmentation can be accomplished using either simple thresholding or machine learning techniques. For cell nuclei, we employed SAM, but Cellpose 2 [38] can be an option. Neuron segmentation can benefit from machine learning technique [39–41].

### Seamless navigation of multiscale vCLEM images using ROI navigation viewer

We developed a specialized viewer for intuitive navigation of image registration results. This tool empowers users to:

1. Visualize 3D relationships: Grasp the spatial context of ROIs within the entire block after LM imaging and registration.
2. Optimize block trimming: Minimize block size by accurately locating ROIs on the ultramicrotome camera view, maximizing section count per grid slot.
3. Estimate cutting depth: Visually assess the proximity of the latest section to the ROI depth.
4. Trace ultrastructure: Seamlessly navigate between ROI FM and ROI TEM maps to explore intricate 3D details, such as axons.

Our method extends the capabilities of Fiji’s Correlia plugin [42] into the third dimension.

### Analysis of cutting depth estimation

We observed a strong linear correlation between the estimated depths of sections and their corresponding feeding values. This linear relationship supports the validity of using image registration to estimate cutting depth. Deviations were noted in deeper regions, potentially due to the decreased contrast in TEM images. For these regions, we utilized Section TEM maps for image registration. The reduced contrast in these deeper Section TEM maps can introduce uncertainties in segmentation and shape determination.

Depth estimation based on image registration contains inherent uncertainty, arising from both the object’s structural complexity and the imaging system’s limitations. For cylindrical objects aligned with the depth axis, especially those with minimal thickness variation, accurate depth estimation is particularly challenging. In images captured with a long depth-of-focus objective lens, objects can appear stretched in the depth dimension within the image stack. Considering these factors, we estimate the depth uncertainty to be approximately 2 micrometers.

### Comparison of the motorized ultramicrotome approach for targeted sectioning

The motorized ultramicrotome leverages x-ray micro-CT data to automatically section blocks to a specific target plane. This automation streamlines the ultramicrotomy process, albeit requiring access to x-ray micro-CT. In contrast, the integrated workflow eliminates this need. Cutting depths of sections were estimated using Section FMs and Section TEM maps, however, this strategy can be applied to sections without fluorescence by using completely Section TEM maps for depth estimation, making our workflow a viable option for targeted vCLEM.

## Materials and methods

### Viral injections and transcardial perfusion fixation

C57BL/6J male mice were used for all experiments. At P1-P2 pups were anaesthetized with Iso-fluorane and intra-cortically injected with 1 ul of a viral solution containing (AAV-EF1A-Gephyrin.FingR-GFP-CCR5TC – Addgene#125692 and AAV-CAG-tdTomato - Addgene# 59462) into layers 2/3 using a Drummond Nanojet microinjector according to the manufacturer’s protocol.

Injected mice were perfused transcardially at P21. A lethal dose of Sodium Pentobarbitone was intraperitoneally injected into pups, prior to perfusion with saline (0.05% in dH2O) via a peristaltic pump (Miniplus 3, Gilson) and 30G x ½ inch needle. Perfusate was switched to formaldehyde (4% in 175 mM sodium phosphate buffer (high osmolarity PB). once the liver had blanched. Perfused mice were decapitated, and brains postfixed in the same solution (10–12 h). Fixed brains were sliced into 100 µm coronal sections using a vibratome (Leica VT10005), and then were stored in regular osmolarity PBS (supplemented with 0.05% NaN3).

### Immunolabelling

Cortical regions containing infected neurons were incubated for 2-h in a blocking buffer (10% goat serum, 5% BSA in high osmolarity PB). They were then incubated overnight with primary antibody against GFP at dilution of 1:1000 in antibody solution (5% goat serum, 1% BSA in high osmolarity PB). Following three 5 minute washes in high osmolarity PB, slices were incubated for 1 h (RT) with nanobody at a dilution of 1:500 (FluoTag-X4 anti-RFP AZDye568; Nanotag Biotechnologies) and secondary antibodies (Anti-Chicken 488, Invitrogen) at a dilution of 1:1000 in antibody solution and rinsed with PBS (4 times 10 min each). Finally, sections were incubated in 1:200 dilution of Lycopersicon Esculentum Lectin (LEL), DyLight649 (Vector Laboratories) and DAPI (1 ug/ml) in PBS for 30 mins at RT.

### Confocal microscopy

Resin embedded tissue blocks were imaged either a Nikon AX-R NSPARC or a SoRA confocal microscope using a custom built imaging chamber. Low magnification reference stacks were imaged using 20 X air objectives using the 405, 488 nm, 543 nm and 650 nm laser lines and collected with the appropriate emission filters. Higher resolution image stacks were obtained using a 60X oil immersion objective.

Ultrathin sections (200-300 nm) were either collected onto glass slides or collected in slot grids and placed onto 35 mm plastic petri dishes containing PBS. They were imaged using an inverted confocal Nikon AX-R NSPARC.

### EM sample preparation

EM sample preparation was performed by following a previous article [25]: Samples were frozen with polyvinylpyrrolidone as a cryoprotectant, in an EM ICE high pressure freezer (Leica). Freeze substitution and resin embedding were performed in an automated AFS2 machine (Leica), using the freeze substitution processor unit. To facilitate the infiltration of Lowicryl HM20 (Polysciences Inc), the temperature was gradually raised to −25°C while increasing the resin concentration in acetone, and the samples were UV polymerized at −25°C through to +25°C.

### Ultramicrotomy

Blocks were sectioned using a UC7 (Leica, Wetzlar, Germany). Section thickness was 200 nm. Feeding values was recorded for cutting each section.

### TEM imaging

#### Section TEM map

The entire section was imaged using a transmission electron microscope (TEM) JEM-1400 Flash (Jeol, Tokyo, Japan) with acceleration voltage of 80 kV, with Limitless Panorama montaging software. Typical montage size was 400 x 400 µm². The pixel size (X/Y) was 340 nm.

#### ROI TEM map

The region round the ROI was imaged by TEM as above, except typical size was 70 × 100 µm². The pixel size (X/Y) was 3.4 nm.

#### TEM tomography

The regions round the ROI were imaged using either a JEM-1400 Flash (Jeol, Tokyo, Japan) TEM with accelerating voltage of 120 kV or a JEM-F200 TEM with a DE16 camera, at 200 kV. Tilt series were acquired using SerialEM [43]. The typical size was 4 x 4 x 0.2 µm^3^ volume. Voxel size was 1.8 nm or 0.94 nm, respectively. The tomograms were reconstructed using IMOD [44,45] software.

### Image preprocessing for registration

Prior to applying for segmentation-based image registration, pixel resolutions of images were matched using Fiji. All the image processing was performed using the NVIDIA A5000 graphics processing unit.

For Section FMs, raw image stacks were MIP projected after drift in Z direction was corrected using Correct 3D Drift plugin [46].

For non-linear image registration between adjacent ROI TEM maps, approximately 30 corresponding points were manually identified and registered using bUnwarpJ plugin [47].

### Segmentation

#### Blood vessels

Image binarization was used in light microscopy images. The threshold for binarization was determined using the maximum entropy method.

#### Cell nuclei

Cell nuclei segmentation algorithms have been actively proposed for both light and TEM images. The SAMJ plugin [48] in Fiji [49] was used. The SAM plugin [50] in napari [51] can also be used. Users identify cell nuclei while viewing light microscopy images and annotate boxes around the identified nuclei. Using the annotated boxes as a prompt, the algorithm segments the cell nuclei.

#### Neurons

Image binarization was used for light microscopy images. The threshold for binarization was determined using the maximum entropy method. Manual segmentation was performed for TEM images.

#### Synapses

Image binarization was used for light microscopy images. The threshold for binarization was determined using the maximum entropy method. Manual segmentation of presynaptic boutons was performed for TEM tomograms.

## Acknowledgements

We would like to thank Pedro Machado for help in development of methods for high pressure freezing and freeze substitution and very useful advice for analysis. We would also like to thank all the members of Burrone lab for useful feedback and comments on the manuscript, and Satoshi Takahashi, Kazuhiro Kido, Erika Kanematsu, Masataka Murakami, and Tetsuya Koike of Nikon Corporation for support. Finally, we would like to thank the Nikon Imaging Centre at Kings College London for help with light microscopy. This research was funded in whole, or in part, by the Wellcome Trust (Grant No. 215508/Z/19/Z to JB). This research was also supported by the European Research Council (692659), the British Heart Foundation (CH/11/3/29051 and RG/16/15/32294), the Fondation Leducq (RA15CVD04).

